# Evolution of copper resistance in the kiwifruit pathogen *Pseudomonas syringae* pv. *actinidiae* through acquisition of integrative conjugative elements and plasmids

**DOI:** 10.1101/070391

**Authors:** Elena Colombi, Christina Straub, Sven Künzel, Matthew D. Templeton, Honour C. McCann, Paul B. Rainey

**Affiliations:** New Zealand Institute for Advanced Study, Massey University, Auckland, New Zealand.; Max Planck Institute for Evolutionary Biology, Plön, Germany.; Plant and Food Research, Auckland, New Zealand.; School of Biological Sciences, University of Auckland, Auckland, New Zealand.; South China Botanical Institute, Chinese Academy of Sciences, Guangzhou, China.; Ecole Supérieure de Physique et de Chimie Industrielles de la Ville de Paris (ESPCI Paris-Tech), PSL Research University, Paris, France.

**Author notes:** Joint senior authors (Honour C. McCann, Paul B. Rainey). Correspondence: Elena Colombi, New Zealand Institute for Advanced Study, Massey University, Private Bag 102 904, Auckland 0745, New Zealand.

## Abstract

**SUMMARY:** Lateral gene transfer can precipitate rapid evolutionary change. In 2010 the global pandemic of kiwifruit canker disease caused by *Pseudomonas syringae* pv. *actinidiae* (*Psa*) reached New Zealand. At the time of introduction, the single clone responsible for the outbreak was sensitive to copper, however, analysis of a sample of isolates taken in 2015 and 2016 showed that a quarter were copper resistant. Genome sequences of seven strains showed that copper resistance – comprising *czc*/*cusABC* and *copABCD* systems – along with resistance to arsenic and cadmium, was acquired via uptake of integrative conjugative elements (ICEs), but also plasmids. Comparative analysis showed ICEs to have a mosaic structure, with one being a tripartite arrangement of two different ICEs and a plasmid that were isolated in 1921 (USA), 1968 (NZ) and 1988 (Japan), from *P. syringae* pathogens of millet, wheat and kiwifruit, respectively. Two of the *Psa* ICEs were nearly identical to two ICEs isolated from kiwifruit leaf colonists prior to the introduction of *Psa* into NZ. Additionally, we show ICE transfer *in vitro* and *in planta*, analyze fitness consequences of ICE carriage, capture the *de novo* formation of novel recombinant ICEs, and explore ICE host-range.

**ORIGINALITY-SIGNIFICANT STATEMENT:** Lateral gene transfer is a major evolutionary force, but its immediacy is often overlooked. Between 2008 and 2010 a single virulent clone of the kiwifruit pathogen *Pseudomonas syringae* pv. *actinidiae* spread to kiwifruit growing regions of the world. After arrival in New Zealand it acquired genetic determinants of copper resistance in the form of integrative conjugative elements and plasmids. Components of these elements are evident in distantly related bacteria from millet (USA, 1921), kiwifruit (Japan, 1988) and wheat (New Zealand, 1968). Additional laboratory experiments capture evidence of the dynamism underpinning the evolution of these elements in real time and further emphasize the potent role that lateral gene transfer plays in microbial evolution.

## INTRODUCTION

Horizontal gene transfer (HGT) is a potent evolutionary process that significantly shapes patterns of diversity in bacterial populations. Horizontally transmissible elements, including plasmids, phages and integrative conjugative elements (ICEs) move genes over broad phylogenetic distances and mediate abrupt changes in niche preferences (Sullivan and Ronson, 1998; Ochman *et al*., 2000; Ochman *et al*., 2005; Guglielmini *et al*., 2011).

ICEs are plasmid-like entities with attributes of temperate phages that disseminate vertically as part of the bacterial chromosome and horizontally by virtue of endogenously encoded machinery for conjugative transfer (Wozniak and Waldor, 2010; Guglielmini *et al*., 2011). Essential genetic modules include those mediating integration, excision, conjugation and regulation of conjugative activity (Mohd-Zain *et al*., 2004; Juhas *et al*., 2007; Roberts and Mullany, 2009). During the process of conjugation ICEs circularize and transfer to new hosts, leaving a copy in the original host genome (Wozniak and Waldor, 2010; Johnson and Grossman, 2015). Conjugation during pathogenesis is often regulated by environmental signals (Lovell *et al*., 2009; Quiroz *et al*., 2011; Vanga *et al*., 2015).

In addition to a set of essential genes, ICEs often harbour “cargo” genes of adaptive significance to their hosts. These include genes affecting biofilm formation, pathogenicity, antibiotic and heavy metal resistance, symbiosis and bacteriocin synthesis (Peters *et al*., 1991; Rauch *et al*., 1992; Ravatn *et al*., 1998; Beaber *et al*., 2002; Drenkard *et al*., 2002; Burrus *et al*., 2006; Ramsay *et al*., 2006; Dimopoulou *et al*., 2007; Kung *et al*., 2010). The genetic information stored in cargo genes varies considerably causing ICEs to range in size from 20 kb to 500 kb (Johnson and Grossman, 2015).

In 2008 a distinct and particularly virulent form of the kiwifruit pathogen *Pseudomonas syringae* pv. *actinidiae* (*Psa*) was identified in Italy. It was subsequently disseminated throughout kiwifruit growing regions of the world causing a global pandemic that reached New Zealand (NZ) in 2010 (Balestra *et al*., 2010; Abelleira *et al*., 2011; Everett *et al*., 2011; Vanneste *et al*., 2011). Genomic analysis showed that although the pandemic was derived from a single clone it acquired a set of distinctive ICEs during the course of its global journey (Mazzaglia *et al*., 2012; Butler *et al*., 2013; McCann *et al*., 2013). The NZ lineage carries *Psa*_NZ13_ICE_eno which harbours a 20 kb “enolase” region that is also found in otherwise divergent *Psa* ICEs (McCann *et al*., 2013; McCann *et al*., 2016).

Copper sprays have long been used in NZ to protect plants from a range of diseases. Since the arrival *Psa* in NZ kiwifruit orchardists have employed copper based products to protect vines. From 2011 an ongoing industry-based programme has been in place to monitor copper resistance. In 2014 evidence was first obtained of *Psa* isolates resistant to copper sulphate. Given that the clone of *Psa* originally introduced into NZ was sensitive to copper and lacked genes encoding copper resistance (McCann *et al*., 2013), detection of copper resistance raised the possibility that the evolution of copper resistance in *Psa* is an evolutionary response to the use of copper-based sprays.

Here we report the phenotypic and genetic basis of copper resistance in NZ isolates of *Psa* and show that its emergence has been fuelled by lateral gene transfer involving ICEs and plasmids. We additionally describe the mosaic structure of ICEs, show the dynamics of ICE transfer both *in vitro* and *in planta*, analyze fitness consequences of ICE carriage, capture the *de novo* formation of novel recombinant ICEs, and explore ICE host-range.

## RESULTS

### Occurrence of copper resistance in *Psa*

*Psa* NZ13, isolated in 2010 and representative of the clone introduced in New Zealand, lacks genes encoding copper resistance (McCann *et al*., 2013) and is unable to grow at copper concentrations in excess of 0.8 mM CuSO_4_. Prior to 2014 no copper resistant or tolerant strains had been reported. However, in 2014, two strains isolated from two different kiwifruit orchards, *Psa* NZ45 and *Psa* NZ47, displayed copper resistance, with a MIC of 1.2 mM CuSO_4_. This finding prompted a small-scale sampling of both copper treated and untreated orchards in 2015/2016 encompassing the area where resistance was first identified. From a sample of 213 strains isolated from seven orchards 59 were found to be copper resistant. Copper resistant isolates were sampled from both copper treated and untreated orchards. Additional copper resistant strains were procured from other kiwifruit-growing regions of New Zealand (Figure S1).

### ICE and plasmid-mediated acquisition of copper resistance in *Psa*

The genome of the focal copper resistant isolate, *Psa* NZ45, is a direct clonal descendant of the isolate originally introduced into NZ (*Psa* NZ13) (McCann *et al*., 2016), but differs in two significant regards. Firstly, the “native” ICE (*Psa*_NZ13_ICE_eno) at *att*-1 (immediately upstream of *clpB*), is located at the second *att* site (*att*-2) immediately downstream of *queC* (Figure 1). Secondly, the genome harbours a new 107 kb ICE (*Psa*_NZ45_ICE_Cu) integrated at the *att*-1 site: *Psa*_NZ45_ICE_Cu carries genes encoding copper resistance (Figures 1 and 2A).

**Figure 1.**
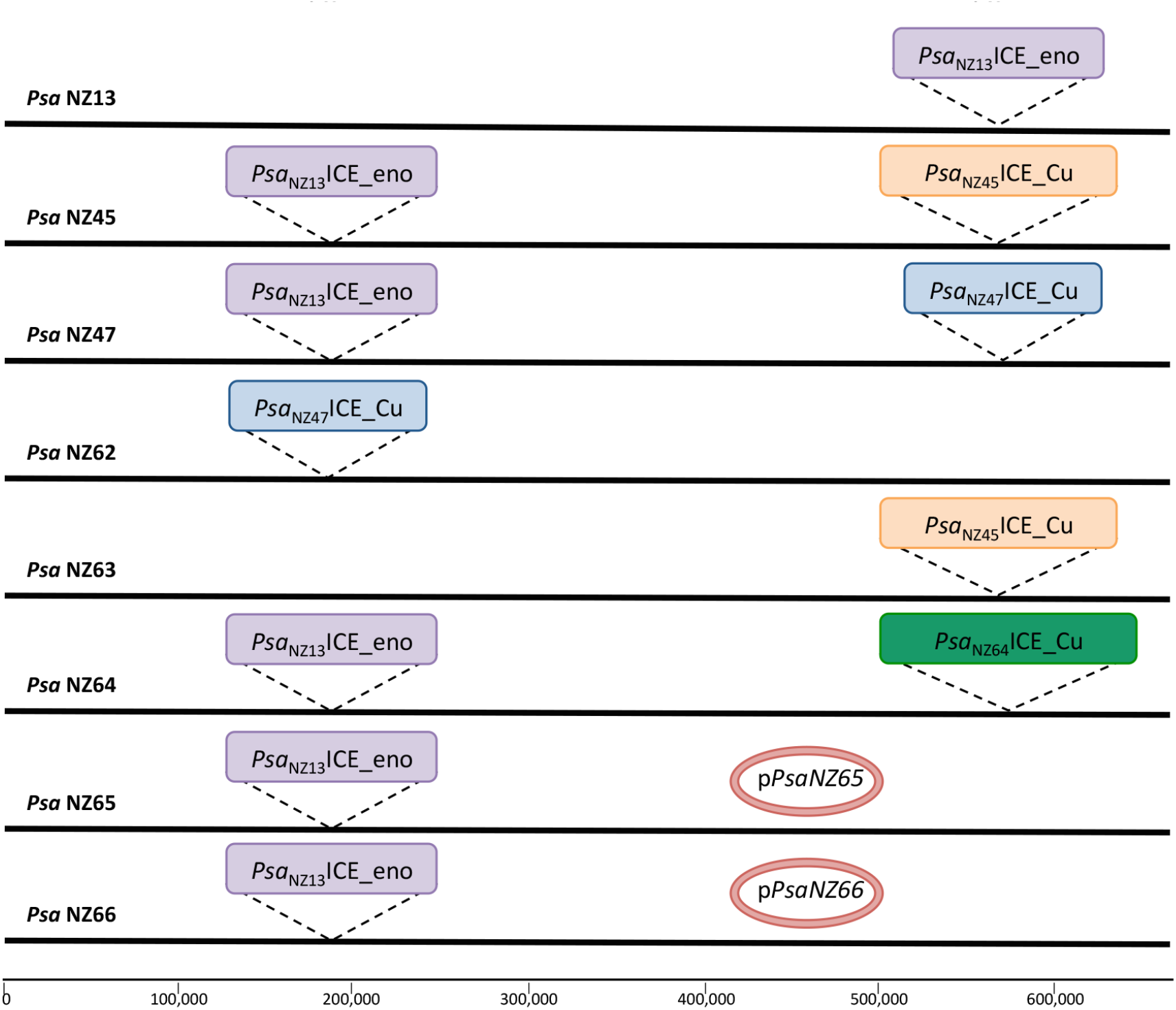
Genomic location of *Psa*ICEs in *Psa* NZ13. In purple the *Psa*_NZ13_ICE_eno (100 kb), in orange *Psa*_NZ45_ICE_Cu (107 kb), in blue the *Psa*_NZ47_ICE_Cu (90 kb), in green the *Psa*_NZ64_ICE_Cu (130 kb), pPsaNZ65 and pPsaNZ66 plasmids are 111 kb. Each island is bounded by 52 bp *att* sequences overlapping tRNALys. In *Psa* NZ13 the *att-1* site is located at 5,534,632 bp, *att-2* at 1,733,972 bp. The figure is not to scale (the entire genome of 6.7 Mbp is indicated a single black line). Both *Psa*_NZ13_ICE_Eno and *Psa*_NZ47_ICE_Cu ICEs were detected in *Psa* NZ47 by sequencing, but analysis of independent colonies from the freezer stock show that Psa_NZ13_ICE_Eno is prone to loss.

**Figure 2.**
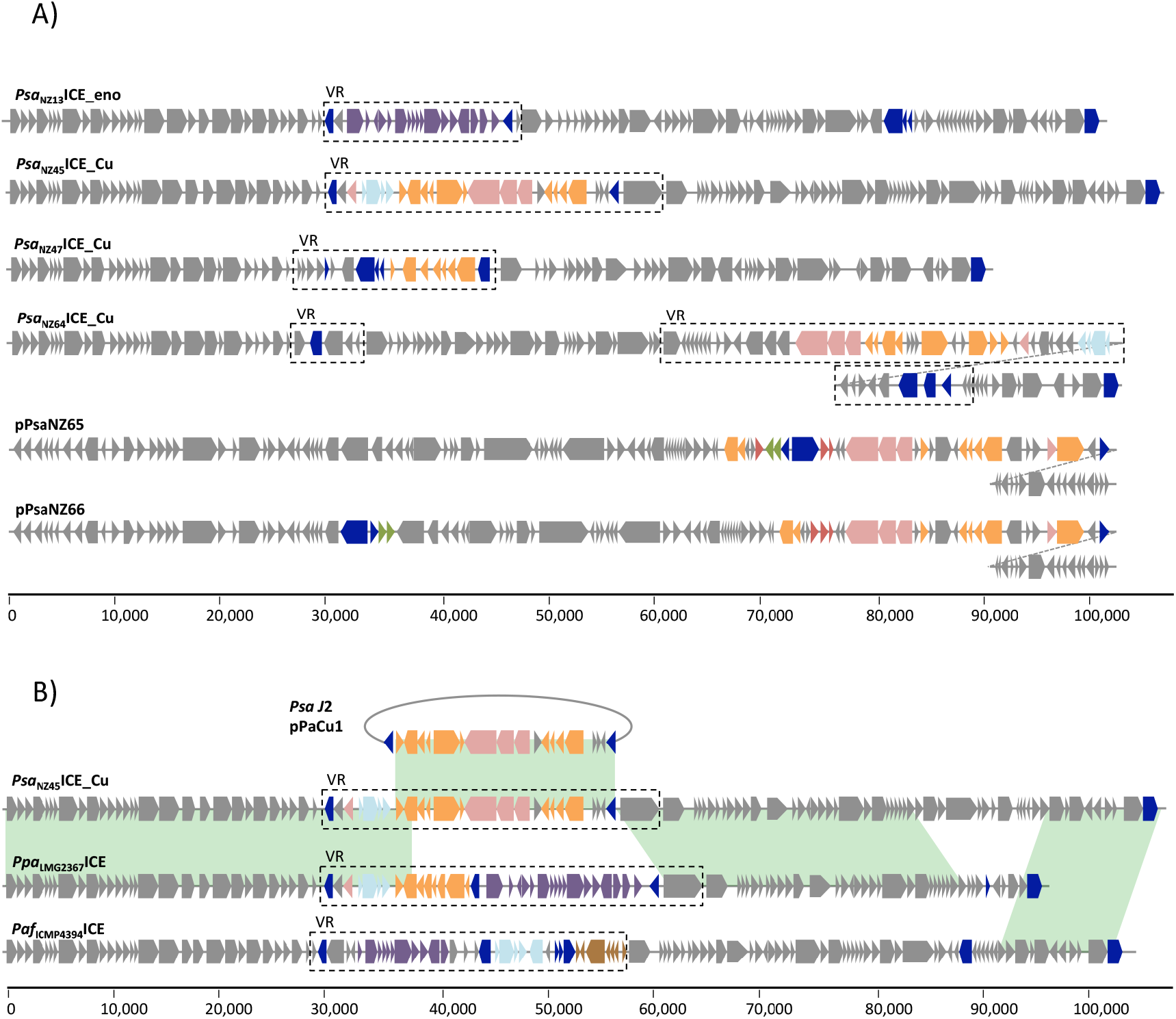
Genetic organization of ICEs and plasmids acquired by *Psa* and mosaicism of *Psa*_NZ45_ICE_Cu. **A)** Blue boxes are mobile genes (transposases or recombinases), purple boxes define the ‘enolase region’, orange boxes depict copper resistance genes, azure boxes are arsenic resistance genes, pink boxes are genes belonging to the *czc*/*cus* system, green boxes are streptomycin resistance genes, red boxes are cation transporter ATPases, brown boxes denote genes encoding mercury resistance. Core “backbone” and other cargo genes are depicted as grey boxes. Dotted diagonal lines indicate continuation of the element.**B)** Areas in green show more than 99% pairwise nucelotide identity. *Psa*_NZ45_ICE_Cu and *Ppa*_LGM2367_ICE share identity both in the first 38 kb and 20 kb downstream of VR. The remaining 20 kb of the *Psa*_NZ45_ICE_Cu VR is almost 37 identical to pPaCu1 (it differs by just 2 SNPs). The last 12.5 kb of *Psa*_NZ45_ICE_Cu is identical to *Paf*_ICMP4394_ICE.

The genomes of six additional copper resistant *Psa* isolates were also sequenced (Table 1) and as with *Psa* NZ45, reads were aligned against the *Psa* NZ13 reference genome (McCann *et al*., 2013; Templeton *et al.,* 2015). All six harbour mobile elements carrying genes encoding copper resistance. The diversity of these elements and genomic location is shown in Figure 1 and their structure is represented in Figure 2A. All isolates are direct clonal descendants of *Psa* NZ13 and thus share an almost identical genome with the exception of the determinants of copper resistance. In *Psa* NZ47 the genes encoding copper resistance are located on a 90 kb ICE (*Psa*_NZ47_ICE_Cu) integrated at the *att-1* site: the native *Psa*_NZ13_ICE_eno is located at the *att-2* site. *Psa* NZ62 carries an ICE identical to that found in *Psa*_NZ47_ICE_Cu (*Psa*_NZ62_ICE_Cu), but is integrated at the *att-2* site; the native ICE (*Psa*_NZ13_ICE_eno) is absent leaving the *att-1* site unoccupied. Isolate *Psa* NZ63 carries *Psa*_NZ45_ICE_Cu integrated at the *att-1* site, but as in *Psa* NZ62, the native *Psa*_NZ13_ICE_eno has been lost. Copper resistance genes in isolate *Psa* NZ64 are also ICE-encoded, but the NZ64 ICE (*Psa*_NZ64_ICE_Cu) is genetically distinct from both *Psa*_NZ47_ICE_Cu and *Psa*_NZ45_ICE_Cu – at 130 kb, it is also the largest. In NZ64, *Psa*_NZ64_ICE_Cu is located at the *att-1* site and the *att-2* site contains the native (*Psa*_NZ13_ICE_eno) ICE. Isolates *Psa* NZ65 and NZ66 both harbour copper resistance genes on a near identical, 120 kb previously undescribed plasmid (pPsaNZ65 and pPsaNZ66, 146 respectively). The only significant difference among the plasmids is the location of a strepomycin resistance-encoding transposon (see below): both isolates have the original *Psa*_NZ13_ICE_eno integrated at the *att-2* site (Figure 1). *Psa* harbouring copper reistance-encoding ICEs have a MIC CuSO_4_ of 1.2 mM while the MIC of 150 plasmid-carrying *Psa* 1.5 mM (Table 1).

**Table 1.**
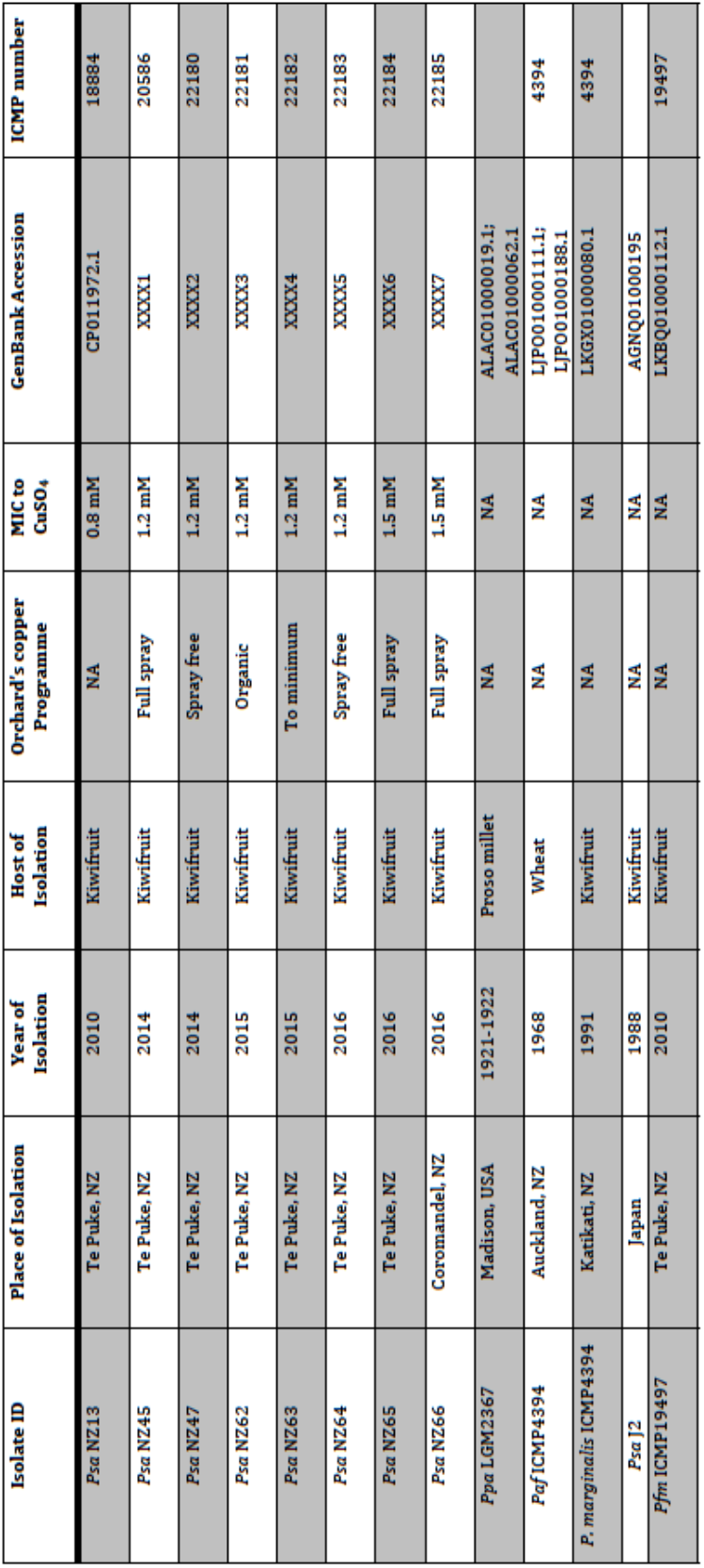
List of genomes used in this study.

That such a small sample of isolates is each unique with regard to the copper resistance-encoding element points to highly dynamic processes shaping their evolution. Such dynamism has been previously noted among enolase-encoding ICEs obtained from a global collection of epidemic *Psa* isolates (McCann *et al*., 2013) and has been observed elsewhere (Burrus *et al*., 2006, Wozniak and Waldorf, 2010). In this study our samples came from a relatively small geographical location. Two ICEs, *Psa*_NZ64_ICE_Cu and *Psa*_NZ47_ICE_Cu were found in different isolates sampled from the same orchard (one year apart), although the two isolates containing near identical plasmids were isolated from orchards located 100 km apart. Two isolates sampled one year apart from the same location (neighboring orchards in Te Puke) carry the same ICE (*Psa*_NZ45_ICE_Cu=*Psa*_NZ63_ICE_Cu; *Psa*_NZ47_ICE_Cu=*Psa*_NZ62_ICE_Cu) (Table 1).

The dynamics of ICE evolution becomes especially evident when placed in the broader context possible by comparisons to ICEs recorded in DNA databases. The core genes of the copper resistance-encoding ICEs from New Zealand *Psa* isolates are syntenous with the core genes of the family of ICEs that includes PPHGI-1 (isolated in 1984 from bean in Ethiopia (Teverson, 1991; Pitman *et al*., 2005) and the three ICEs previously described from *Psa* found in New Zealand (2010), Italy (2008) and China (2010) (McCann *et al*., 2013; Butler *et al*., 2013; E. Colombi, unpublished). *Psa*_NZ45_ICE_Cu is a mosaic of DNA from two known ICEs and a plasmid. It shares regions of near perfect identity (over 66 kb) with ICEs present in the otherwise divergent host genomes of *P. syringae pv. panici* (*Ppa*, LGM2367) isolated from proso millet in Madison (USA) in the 1920s (over the first 38 kb it differs by just 12 SNPs, and one 144 bp insertion), *P. syringae* pv. *atrofasciens* (*Paf*, ICMP4394) isolated in NZ in 1968 from wheat, and a 70.5 kb plasmid present in a non-pandemic *Psa* strain (J2), isolated in Japan in 1988 (Figure 2B).

Interestingly, two of the ICEs described here have also been found in non-*Psa Pseudomonas* isolated from kiwifruit leaves. *Psa*_NZ47_ICE_Cu shows 99.7% pairwise nucleotide identity with an ICE found in *P. marginalis* ICMP 11289 isolated in 1991 from *A. deliciosa* in Katikati (New Zealand). *Psa*_NZ64_ICE_Cu is almost identical (99.5% nucleotide pairwise identity) to an ICE from *P. syringae* pv. *actinidifoliorum* (*Pfm*) ICMP19497, isolated from kiwifruit in 2010 in Te Puke (New Zealand) (Visnovsky *et al*., 2016) (Table 1). Additionally, a 48 kb segment of coding copper resistance genes *Psa*_NZ64_ICE_Cu shares 99.3% nucleotide pairwise identity with a locus found in *P. azotoformans* strain S4 (Fang *et al*., 2016), which was isolated from soil in 2014 in Lijiang (China). However, the locus from *P. azotoformans* is not associated with an ICE.

### Genetic determinants of copper resistance

Copper resistance is typically conferred by operons encoding either copper efflux (*cusABC*) and / or sequestration (*copABCD*) systems, both of which can be under the regulation of the *copRS* / *cusRS* two-component regulatory system (Bondarczuk and Piotrowska-Seget, 2013). ICEs identified in *Psa* isolates harbour operons encoding examples of both resistance mechanisms (and regulators), plus genetic determinants of resistance to other metal ions. In each instance the resistance genes are located within “variable regions” (VR) of ICEs into which cargo genes preferentially integrate (Figure 2A). Delineation of these variable regions comes from detailed analysis of 32 unique ICEs from the *Pph* 1302A and *Psa* families that will be published elsewhere (E. Colombi, unpublished). Overall there are notable similarities and differences in the organization of the variable regions.

As shown in Figure 2B, the first 38 kb of *Psa*_NZ45_ICE_Cu is almost identical (99.7% identical at the nucleotide level) to *Ppa*_LGM2367_ICE. This region spans the core genes, but extends ∼8.2 kb into the variable cargo genes with just two SNPs distinguishing the two ICEs (across the 8.2 kb variable region). Encoded within this region is an integrase, arsenic resistance genes (*arsRBCH*), a gene implicated in cadmium and cobalt resistance (*czcD*) and the *copRS* regulatory system. Partway through *copS* the two ICEs diverge at a recombination breakpoint with the downstream variable region from *Psa*_NZ45_ICE_Cu being homologous to a set of copper resistance genes found on plasmid pPaCu1 from the divergent (non-pandemic) Japanese isolate of *Psa* (J2) (Nakajima *et al*., 2002). This region comprises a putative copper transporting ATPase encoded by *copG* (Gutiérrez-Barranquero *et al*., 2013)*, cusABC* genes involved in the detoxification of monovalent cations, including copper and silver (Mergeay *et al*., 2003; Rensing and Grass, 2003) and *copABCD* (Figure 2B). The last 4 kb of the variable region of *Psa*_NZ45_ICE_Cu shares almost complete identity with *Ppa*_LGM2367_ICE (Figure 2B).

Detail of the diversity of copper resistance (and related metal resistance) genes is shown in Figure 3. All elements (ICEs and plasmids) harbour the *copRS* regulatory system and, with the exception of *Psa*_NZ47_ICE_Cu, all carry both *cusABC* and *copABCD*, although their organization varies. For example, while *copABCD* is typical, in *Psa*_NZ64_ICE_Cu *copAB* and *copCD* are organized as two separate operons (Figure 3). The putative copper ABC transport system encoded by *copG* is a common feature, and determinants of arsenic resistance are present in both *Psa*_NZ45_ICE_Cu and *Psa*_NZ64_ICE_Cu. The putative cadmium and related metal resistance gene, *czcD* is also present on these two ICEs. As noted above, a transposon carrying determinants of streptomycin resistance (*strAB*) is present on plasmids pPsaNZ65 and pPsaNZ66. The transposon is of the Tn*3* family and the cassette bears identity to streptomycin resistance carrying transposons found in *P. syringae* pv. *syringae* B728a (Feil *et al*., 2005), but also on plasmid pMRVIM0713 from *Pseudomonas aeruginosa* strain MRSN17623 (GenBank: KP975076.1), plasmid pPMK1-C from *Klebsiella pneumoniae* strain PMK1 (Stoesser *et al*., 2014), and plasmid pTi carried by *Agrobacterium tumefaciens* LBA4213 (Ach5) (GenBank: CP007228.1).

**Figure 3.**
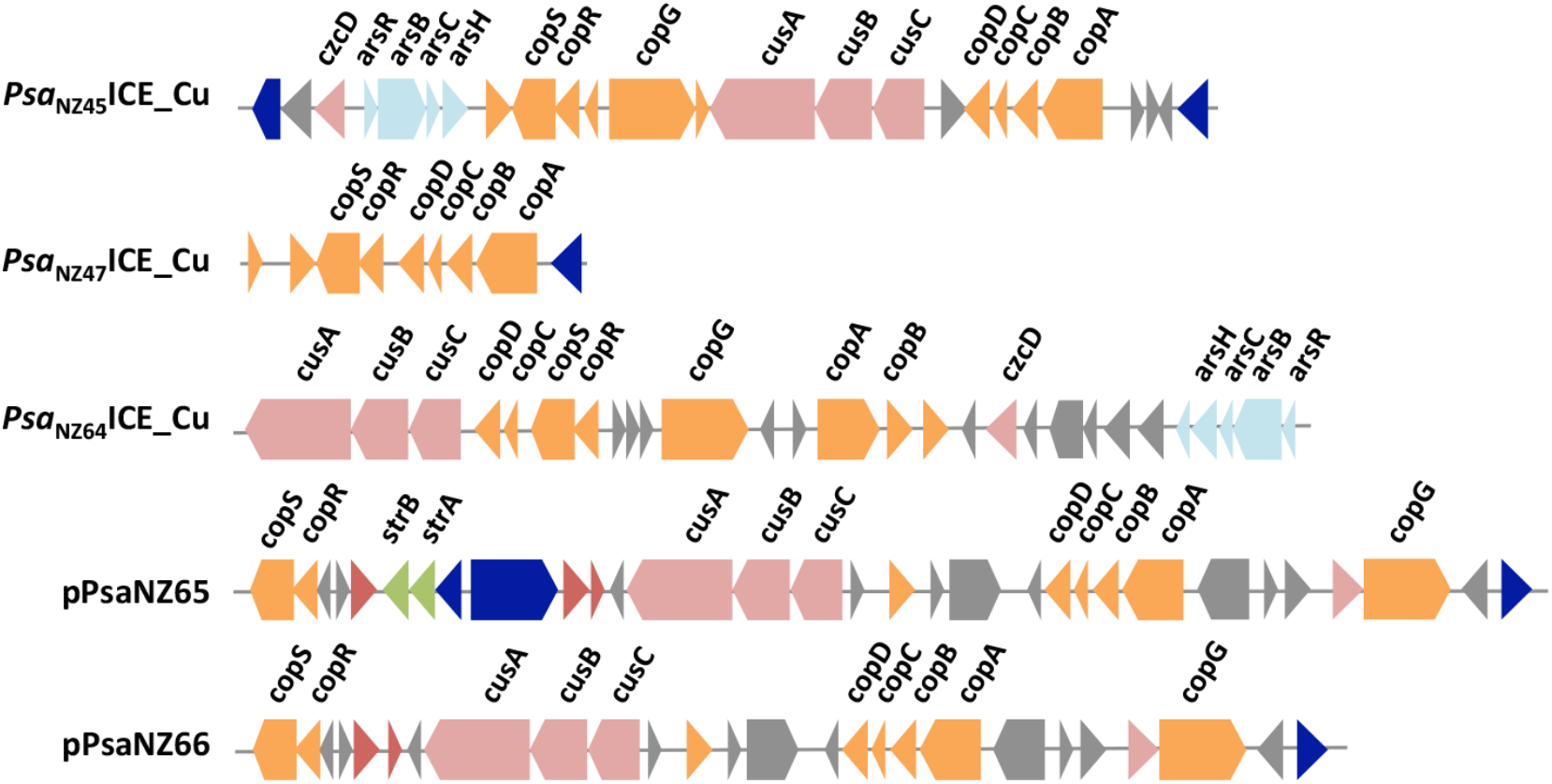
Genetic organization of metal resistance loci. Blue boxes are mobile genes (transposases or recombinases), orange boxes depict copper resistance genes, azure boxes are arsenic resistance genes, pink boxes are genes belonging to the *czc/cus* system, green boxes are streptomycin resistance genes and other genes are depicted as grey boxes.

At the level of the operons determining copper resistance there is marked genetic diversity, however, with the exception of CopR, there is relatively little evidence of within operon recombination. The CusABC system is carried on pPsaNZ65 and pPsaNZ66 (but these are identical) and the ICEs *Psa*_NZ45_ICE_Cu and *Psa*_NZ64_ICE_Cu: CusA, CusB and CusC show 75.8%, 50.0% and 44.8% pairwise amino acid identity, respectively; phylogenetic trees based on protein sequences show congruence (Figure S2). The CopABCD system is present on *Psa*_NZ45_ICE_Cu, *Psa*_NZ47_ICE_Cu, *Psa*_NZ64_ICE_Cu (but CopAB and CopCD are in different locations (Figure 3)) and plasmid pPsaNZ65 (and pPsaNZ66): CopA, CopB, CopC and CopD show 76.4%, 63.1%, 79.1% and 60.8% pairwise amino acid identity, respectively. With the exception of CopC (where bootstrap support is low) phylogenetic trees for each protein show the same overall arrangement (Figure S3). The two component regulatory system *copRS* is also present on each of the elements with the amino acid sequences of CopR showing 84.3% and those of 11 CopS 63.0% pairwise amino acid identity. Phylogenetic trees show CopS from *Psa*_NZ64_ICE_Cu to be the most divergent, and those from *Psa*_NZ45_ICE_Cu and *Psa*_NZ47_ICE_Cu being most similar: CopR shows the same phylogenetic arrangement, however, CopR sequences from *Psa*_NZ45_ICE_Cu and *Psa*_NZ47_ICE_Cu are identical at the protein level suggesting a recent recombination event (Figure S4).

### *Psa*_NZ45_ICE_Cu imposes no detectable fitness cost and confers a selective advantage *in vitro* in the presence of copper

To determine whether ICE carriage confers a fitness cost, we took advantage of the fact that *Psa* NZ13 and *Psa* NZ45 are essentially isogenic, with the exception of the additional ICE in *Psa* NZ45 (*Psa*_NZ45_ICE_Cu). Each strain was grown alone and density of cells monitored over a 72 hour period with samples taken every 24 hours. In the absence of copper sulphate, no difference in cell density was detected; however, in the presence of 0.5 and 0.8 mM CuSO_4_ the density of *Psa* NZ13 was reduced (Figure 4A). There is thus no apparent fitness cost associated with carriage of *Psa*_NZ45_ICE_Cu in the absence of copper sulphate, but there is a fitness advantage in copper-containing environments.

**Figure 4.**
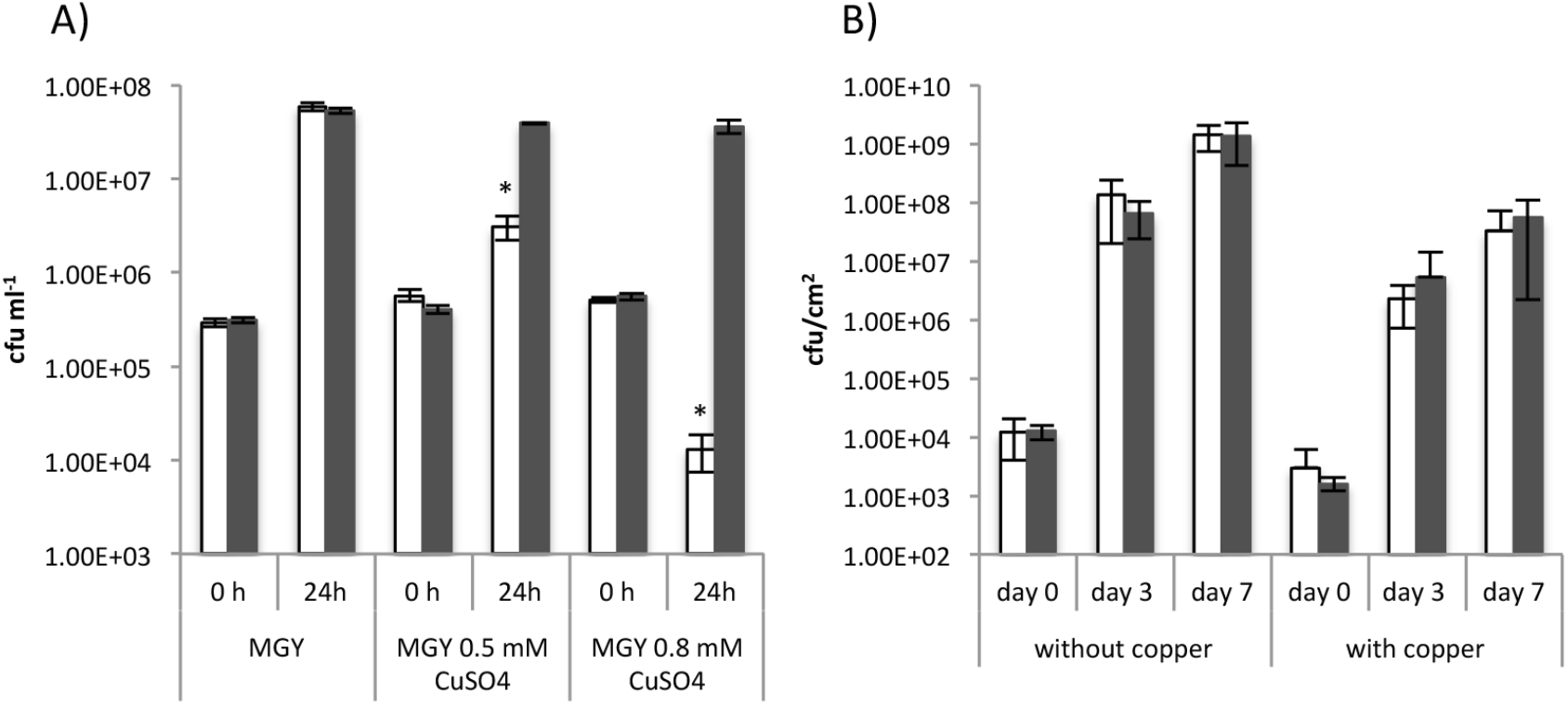
Effect of copper ions on growth of *Psa* NZ13 and *Psa* NZ45. **A)** *Psa* NZ13 (white bars) and *Psa* NZ45 (grey bars) were grown for 24 h in shaken MGY culture and MGY supplemented with 0.5mM and 0.8 mM CuSO_4_. Data are means and standard deviation of three independent cultures. *indicates significant difference *P*<0.05 (one tailed *t*-test)). **B)** The single growth of *Psa* NZ13 (white bars) and *Psa* NZ45 (grey bars) was assessed endophytically on leaves of the kiwifruit cultivar Hort16A. Data are means and standard deviation of five replicates. The copper product Nordox75 (0.375g L-1) was sprayed adaxially and abaxially until run off. Data are means and standard deviation of 5 replicates. One tailed *t-*test showed no statistical difference in growth between of *Psa* NZ13 and NZ45 in absence or presence of copper.

Although carriage of *Psa*_NZ45_ICE_Cu appeared not to affect the growth of *Psa* NZ45 in the absence of copper, a more precise measure of fitness was sought by performing competition experiments in which *Psa* NZ13 and *Psa* NZ45 were co-cultured. For this experiment *Psa* NZ13 was marked with a kanamycin resistance cassette so that it could be distinguished from kanamycin sensitive, copper resistant *Psa* NZ45. Over a 24 hour period where the two strains (founded at equal density) competed for the same resources (in shaken MGY 12 medium without copper sulphate), the fitness of *Psa* NZ45 was not significantly different to *Psa* NZ13 (1.07 ± 0.04; mean and SEM from 3 independent experiments, each comprised of 3 replicates), indicating no significant detectable cost of carriage of *Psa*_NZ45_ICE_Cu.

Given that the mechanism of copper resistance in *Psa* NZ45 – based upon *copABCD* – likely involves sequestration of copper ions we considered the possibility that this isolate might confer cross protection to non-copper resistant isolates, such as *Psa* NZ13. To this end we performed co-culture experiments at sub-inhibitory and inhibitory copper sulphate concentrations. Growth of *Psa* NZ13 at sub-inhibitory concentrations of copper sulphate was significantly impaired by the presence of *Psa* NZ45 and this was especially evident at 48 and 72 hours (Figure S5). At the inhibitory copper sulphate concentration, *Psa* NZ13 appeared to benefit from the presence of *Psa* NZ45. Again, this was most evident at 48 and 72 hours (Figure S5).

### *Psa*_NZ45_ICE_Cu imposes no detectable fitness cost and confers no selective advantage *in planta*

Cost and benefit of carrying *Psa*_NZ45_ICE_Cu was also evaluated during endophytic colonization of kiwifruit leaves. No significant difference was observed in growth of singly-inoculated *Psa* NZ13 and NZ45 (dip inoculation was used to found colonization). Spray application of a commonly used commercial copper-based treatment (Nordox75 (0.375 g L^-1^)) subsequent to dip inoculation resulted in a reduction of bacterial density of both strains and the presence of *Psa*_NZ45_ICE_Cu in *Psa* NZ45 did not confer any advantage *in planta* (Figure 4B). Co-cultivation competition assays in the presence or absence of Nordox75 confirmed carriage of *Psa*_NZ45_ICE_Cu imposes no significant fitness cost or advantage during endophytic growth: fitness of NZ45 relative to NZ13 was 1.00 ± 0.02 and 1.07 ± 0.03 at day 3 and 7, respectively; fitness of NZ45 relative to NZ13 in the presence of 0.375 g L^-1^ Nordox was 1.15 ± 0.04 and 0.97 ± 0.09 at day 3 and 7, respectively (data are means and SEM from 3 independent experiments, each comprised of 5 replicates; significance was calculated by one sample *t*-test).

### *Psa*_NZ45_ICE_Cu transfer dynamics *in vitro* and *in planta*

Acquisition of *Psa*_NZ45_ICE_Cu (and related ICEs) by *Psa* NZ13 suggests that the element is active and capable of self-transmission. If so, then it is possible that transfer may have occurred during the course of the co-cultivation experiments used to determine cost of ICE carriage. To determine whether this had happened samples from the mixtures were plated on MGY medium containing both kanamycin and copper sulphate. Copper resistant, kanamycin resistant transconjugants were detected both *in vitro* and *in planta.* This means that a fraction of *Psa* NZ13 strains acquired *Psa*_NZ45_ICE_Cu. These transconjugants marginally inflate the counts of *Psa* NZ45, however, the number of transconjugants (see below) was several orders of magnitude less that *Psa* NZ13, thus having no appreciable effect on the measures of relative fitness.

At 24 hours in shaken MGY broth transconjugants were present at a frequency of 5.04 ± 2.25 × 10^−3^ per recipient cell (mean and SEM from 3 independent experiments, each comprised of 3 replicates). Analysis of samples from *in planta* experiments showed that at 3 days, transconjugants were present at a frequency of 2.05 ± 0.63 × 10^−2^ per recipient cell (mean and SEM from 3 independent experiments, each comprised of 5 replicates). On plants in the presence of Nordox (0.375 g L^−1^) the frequency of transconjugants was 9.37 ± 1.56 × 10^−2^ per recipient three days after inoculation (mean and SEM from 3 independent experiments, each comprised of 5 replicates). Transfer was also observed in M9 agar, on M9 agar supplemented with 0.5mM CuSO_4_, on M9 agar supplemented with a macerate of Hort16A fruit with transconjugants present (at 48 hours) at a frequency of 2.16 ± 0.9 × 10^−5^, 1.11 ± 0.4 × 10^−5^ and 1.98 ± 0.8 × 10^−5^ per recipient cell, respectively.

To explore the dynamics of transfer *in vitro*, samples from shaken MGY cultures were taken hourly, for six hours, and then at 24 hours. The data, presented in Figure 5, show acquisition of *Psa*_NZ45_ICE_Cu by *Psa* NZ13 within one hour of the mating mix being established (approximately 1 recipient per 10^5^ recipient cells). The frequency was relatively invariant over the subsequent six hour period, but rose to approximately 1 recipient in 10^3^ cells at 24 hours.

**Figure 5.**
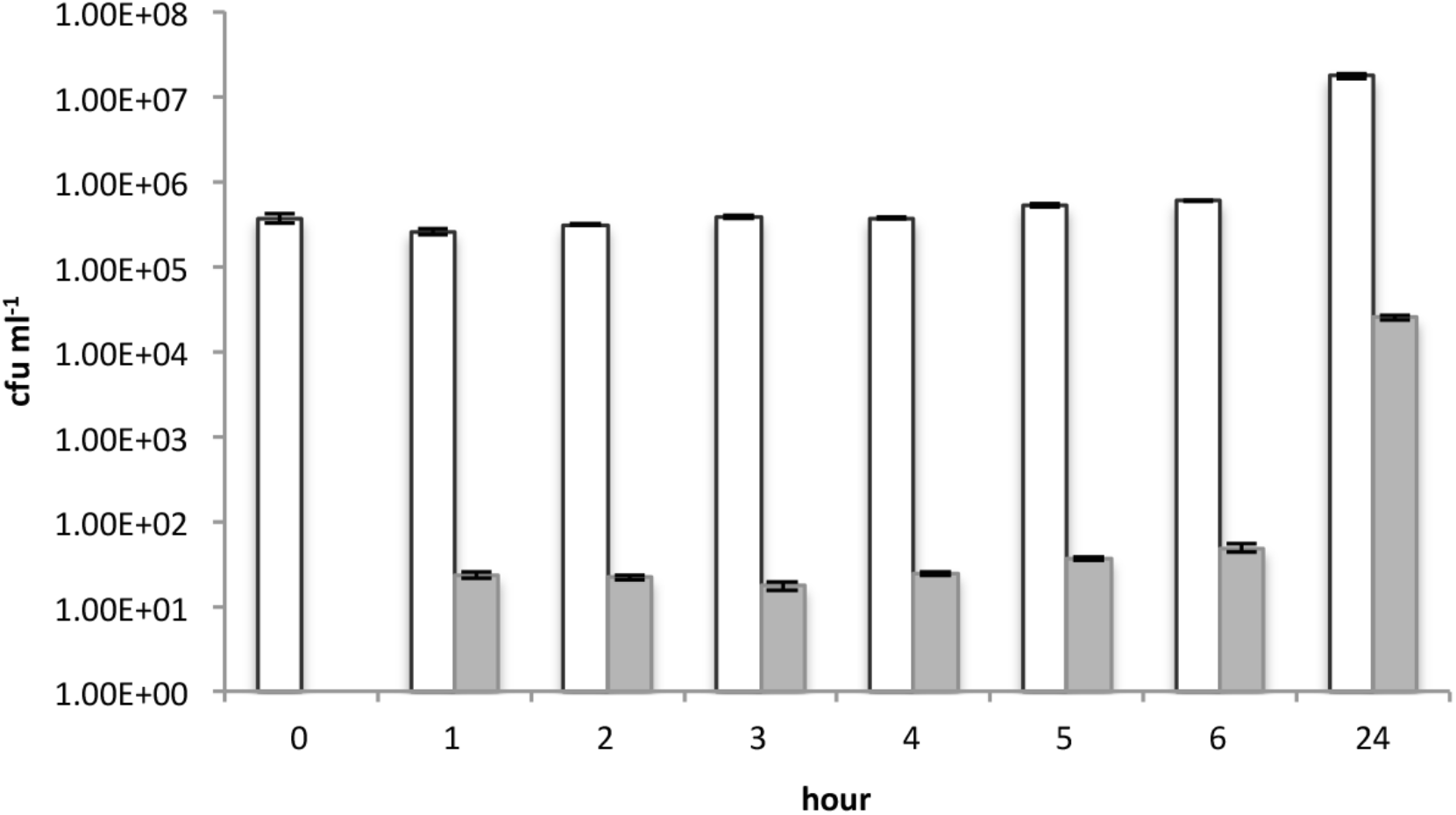
*In vitro* transfer of *Psa*_NZ45_ICE_Cu from *Psa* NZ45 to *Psa* NZ13. Colony forming units of the recipient *Psa* NZ13 (white bars) and *Psa* NZ13 carrying *Psa*_NZ45_ICE_Cu (transconjugants, grey bars) was monitored during co-cultivation. Data are means and standard deviation of 3 independent cultures

Detection of ICE transfer just one hour after mixing donor and recipient cells promoted a further experiment in which transconjugants were assayed at 10 minute intervals. From three independent experiments, each with five replicates, transconjugants were detected at 30 mins (approximately 4 × 10^−7^ transconjugants per recipient cell).

Analysis of co-cultivation experiments from kiwifruit leaves showed evidence that *Psa*_NZ45_ICE_Cu also transferred *in planta.* The frequency of transconjugants at day 3 and day 7 was approximately 1 per 50 recipient cells and the frequency of transconjugants was not affected by changes in the initial founding ratios of donor and recipient cells (Table S1). Overall, the frequency of transconjugants was approximately three orders of magnitude greater *in planta* than *in vitro.*

### ICE displacement and recombination

To check the genetic composition of transconjugants and to investigate whether *Psa*_NZ45_ICE_Cu integration in recipient cells occurred at the *att-1* or *att-2* site, a set of primers were designed to identify the location of ICE integration in the *Psa* NZ13 genome (Table S2). 11 independently generated transconjugants from shaken MGY culture were screened. As expected, successful amplification of primers annealing to *copABCD* in *Psa*_NZ45_ICE_Cu was observed in all transconjugants, while amplification of the *enolase* gene primers (indicative of the presence of the native *Psa*_NZ13_ICE_eno) occurred only in *Psa* NZ13 and *Psa* NZ45 (Figure S6). However, in two transconjugants only the *IntPsaNZ13-att-1* primer pair resulted in amplification, suggesting that recombination between *Psa*_NZ45_ICE_Cu and *Psa*_NZ13_ICE_eno had occurred (Figure S7). Genome sequencing of one of these transconjugants revealed a recombination event inside the variable region of the ICE that produced a chimeric ICE identical to *Psa*_NZ45_ICE_Cu up to and including the CuR operon, with the remainder identical to the downstream segment of *Psa*_NZ13_ICE_eno (Figure 6).

**Figure 6.**
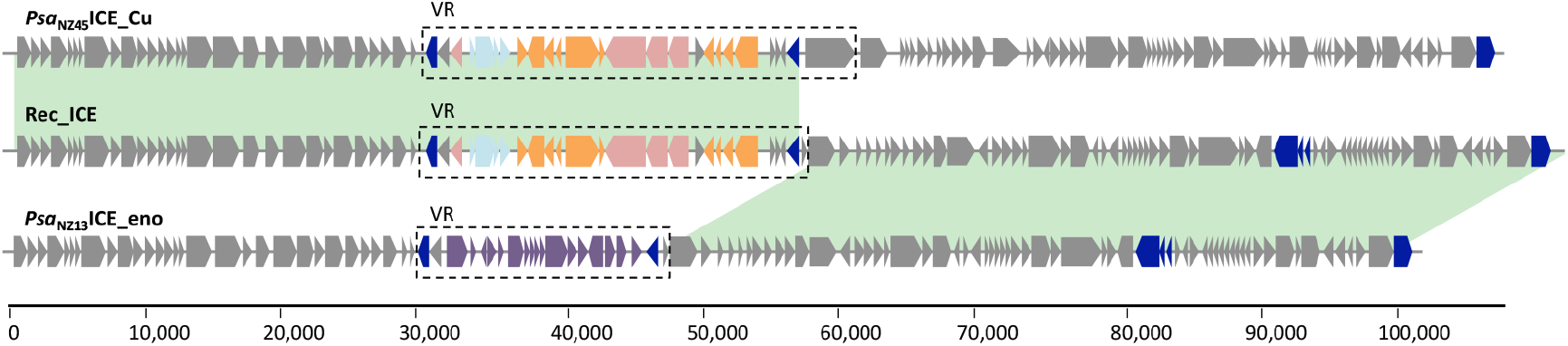
Structure and mosaicism of the recombinant ICE (Rec_ICE) in transconjugant *Psa* NZ13. Areas highlighted in green show 100% pairwise identity. The recombination break point is inside the variable region (VR). Blue boxes are mobile genes (transposases or recombinases), purple boxes define the ‘enolase region’ (McCann *et al*. 2013), orange boxes depict copper resistance genes, azure boxes are arsenic resistance genes and pink boxes are genes belonging to the *czc*/*cus* system. Core “backbone” and other cargo genes are depicted as grey boxes.

### *Psa*_NZ45_ICE_Cu can be transferred to a range of *P. syringae* strains

The host range of the *Psa*_NZ45_ICE_Cu was characterised using a panel of nine different *Pseudomonas* strains as recipients, representing the diversity of *P. syringae* and the genus more broadly. Transfer of *Psa*_NZ45_ICE_Cu to *Psa* J31, *Pfm* NZ9 and *P. syringae* pv. *phaseolicola* (*Pph*) 1448a (on M9 agar plates) was observed with the frequency of transconjugants per recipient cell being 7.64 ± 1.7 × 10^−6^, 7.74. ± 2.5 × 10^−7^ and 1.23 ± 0.2 × 10^−4^, respectively. No transconjugants were detected for *P. aeruginosa* PAO1, *P. fluorescens* SBW25, *P. syringae* pv. *tomato* DC3000 or *Psa* K28, despite the fact that these three strains have both *att* sites.

## DISCUSSION

The importance and impact of lateral gene transfer on the evolution of microbial populations has long been recognized (Sullivan *et al*., 1995; Lilley and Bailey, 1997; Ochman *et al*., 2000; Ochman *et al*., 2005; Wozniak and Waldor, 2010; Polz *et al*., 2013). Here we have captured the real time evolution of copper resistance in a plant pathogen, in an agricultural setting, and shown that movement of copper resistance genes occurs primarily via ICEs. The strains subject to genomic analysis provide a glimpse of just how dynamic evolution fuelled by ICEs can be. Of the seven copper resistant *Psa* isolates analyzed, five contain copper resistance-encoding ICEs – three unique ICEs in total – with variable placement within the *Psa* genome, including movement and instability of the native ICE (*Psa*_NZ13_ICE_eno). Further evidence of dynamism comes from *in vitro* and *in planta* studies, which show not only transfer to isogenic *Psa* and unrelated *P. syringae* strains, but also the ready formation of chimeras between *Psa*_NZ45_ICE_Cu and *Psa*_NZ13_ICE_eno. Mosaicism of ICEs has been reported elsewhere and is often promoted by the presence of tandem copies (Garriss *et al*., 2009; Wozniak and Waldor, 2010). The ease with which ICEs move between strains and capacity for intra-ICE recombination emphasizes the futility of drawing conclusions on strain phylogeny based on ICE phylogeny (McCann *et al*., 2013), but also the impossibility of understanding ICE evolution based on the phylogeny of ICEs themselves.

Evidence of the formation of chimeric ICEs extends beyond the ICEs studied here. *Psa*_NZ45_ICE_Cu is a recombinant of two previously reported ICEs and a plasmid: most surprising is the fact that the recombinant components are derived from elements isolated from three geographic regions (USA, Japan and New Zealand) from three different plants (millet, kiwifruit and wheat) and spanning almost 100 years. Additionally, two of the copper resistance-encoding ICEs found in *Psa* (*Psa*_NZ47_ICE_Cu and *Psa*_NZ64_ICE_Cu) have been reported in other kiwifruit leaf colonizing organisms emphasizing the ease by which self-transmissible elements can move between members of a single community. Clearly the potency of evolution fuelled by ICEs with the *P. syringae* complex is remarkable, with impacts likely extending well beyond that inferred from the analysis of genome sequences (Fondi *et al*., 2016).

Evidence of the spectrum and dynamic of transfer inferred from the genomic analysis of natural isolates is bolstered by demonstration of the *in vitro* and *in planta* transfer of *Psa*_NZ45_ICE_Cu. The fact that *Psa*_NZ45_ICE_Cu can be detected in a recipient strain just 30 minutes after mixing with a donor strain (in shaken broth culture) points to an as yet undetermined proficiency for transfer and possible regulatory mechanism. At the same time, the frequency of transconjugants *in planta* are several orders of magnitude greater than *in vitro* suggesting even greater potential (perhaps regulated) for transfer in the natural environment.

The selective causes underpinning the evolution of copper resistance in *Psa* is uncertain and to date copper resistance is not known to have evolved outside of New Zealand. While it is tempting to blame use of copper sprays by orchardists, it is possible that the evolution of copper resistant *Psa* is a more general response to copper levels in New Zealand soils combined with long-term use of copper-based sprays in New Zealand agriculture (Morgan and Taylor, 2004). Support for this stems from the fact that *Psa*_NZ47_ICE_Cu shows almost perfect identify with an ICE found in *P. marginalis* (ICMP 11289) from kiwifruit isolated in 1991 (in New Zealand). In addition, copper resistance-encoding ICEs were found in both copper treated, and untreated orchards. There is need to understand further the population ecology of copper-resistance ICEs at regional, national and global scales and the selective causes for their maintenance and spread (Staehlin *et al*., 2016).

The impact of the copper resistance-encoding ICEs on fitness *in planta* –in the presence of copper sprays – appears to be minimal. While this is heartening news from the perspective of control of the pathogen, there are at least three reasons to treat this result with caution. Firstly, it is difficult to accurately assess fitness *in planta* and it is possible that our measures underestimate the contribution of copper resistance to growth in the presence of copper: even a 1% increase in fitness over 24 hours, which is beyond experimental capacity to detect, can have significant long-term consequences. Secondly, the presence of copper resistance genes means opportunity for levels of resistance to increase through, for example, promoter mutations that increase levels of transcription of resistance determinants, or through acquisition of additional copper resistance-encoding genes. Thirdly, and perhaps most significantly, is the fact that the copper resistance-encoding ICEs confer no measurable fitness cost even in the absence of copper. This suggests that these elements will not be readily lost from *Psa* populations even if copper-based sprays were eliminated (Andersson and Hughes, 2010; Neale *et al*., 2016). That some strains of the global pandemic now contain two ICEs gives reason to suspect elevated evolutionary potential among these isolates.

While the focus of our investigation has been copper resistance, the ICEs reported here carry a cargo of additional genes, some of which are implicated in resistance to other metals. In some instances the cargo genes have no similarity to genes of known function (grey boxes in Figure 2A). ICEs and similar laterally transferred elements provide opportunity for genes unrelated to copper resistance, for example gene connected to virulence, to hitchhike and rapidly spread. In this regard the two plasmids characterized here are of interest: both carry determinants of streptomycin resistance – an antibiotic that is also sprayed on New Zealand kiwifruit orchards in order to control *Psa.* The potential for hitchhiking has been previously noted in the context of antibiotic resistance-encoding plasmids (Gullberg *et al*., 2014).

Recognition of ICEs along with their potential to change the course of microbial evolution extends less than twenty years (Wozniak and Waldor, 2010). While it might be argued that this potential is no different from that long realized via conjugative plasmids, or phage (Ochman *et al*., 2000), ICEs, being a composite of both, seem to have an edge. Unlike conjugative plasmids that rarely integrate into the host genome, ICEs integrate as a matter of course and are largely immune to segregational loss; additionally, fitness consequences as a result of carriage are likely to be minimal. Unlike temperate phages, ICEs do not kill the host upon transfer, but they can nonetheless mediate transfer upon encountering transfer proficient conditions. Having control over both vertical and horizontal modes of transmission, while minimizing costs for host cells, marks these elements as especially potent vehicles of microbial evolution.

## EXPERIMENTAL PROCEDURES

### Strains and culture condition

All *Pseudomonas* strains were cultured in King’s B medium at 28˚C, *E. coli* was cultured in Luria Bertani medium at 37˚C. All liquid overnight cultures were shaken at 250 rpm. Both kanamycin and nitrofurantoin were used at 50 µg mL^−1^.

### DNA extraction and genome sequencing

For genome sequencing, DNA samples were extracted using the Promega Wizard® Genomic DNA Purification Kit following the recommended protocol. *Psa* NZ45 was sequenced using the PacBio platform, the remainder were sequenced using the Illumina HiSeq platform. Sequences are deposited at NCBI with the following accession numbers: XXXX1, XXXX2 etc (right number of accession numbers).

### Genomic reconstruction of ICEs

ICEs identified in genome sequences were used as query sequences for BLAST searches of the NCBI WGS database (http://blast.ncbi.nlm.nih.gov/Blast.cgi.) Contigs were subsequently downloaded and where ICEs were represented by two contigs they were concatenated in Geneious http://www.geneious.com, Kearse et al., 2012), Kearse *et al*., 2012). Concatenation was only required in two instances. ICEs were annotated using the RAST server http://rast.nmpdr.org, Aziz *et al*., 2008) and manually curated. Alignments were performed using Geneious.

### *Psa* isolation from kiwifruit orchards

One cm^2^ kiwifruit leaf disks were macerated in 200 µl 10mM MgCl_2_. The macerate was plated on *Pseudomonas* selective media amended with cetrimide, fucidin and cephalosporin (Oxoid) and incubated at 28˚C for 3 days. *Psa* was identified using either diagnostic PCR or LAMP assays (Rees-George *et al*., 2010, Ruinelli *et al*., 2016).

### Copper resistance assays

Copper resistance was evaluated by determining the minimal concentration of copper that inhibited growth (minimal inhibitory concentration, MIC) on mannitol-glutamate yeast extract medium (MGY) plates supplemented with CuSO_4_·5H_2_O (Bender and Cooksey, 1986, Cha and Cooksey 1991). *Psa* strains were considered resistant when their MIC exceeded 0.8 mM CuSO_4_.

### Mutant development

A Tn*5* transposon was used to generate kanamycin resistant (*kanR*) strains. *E. coli* S17-1 Tn*5hah Sgid1* (Zhang *et al*., 2015) was used as donor and *E. coli* pK2013 (Ditta *et al.,* 1980) as helper. Helper, donor and recipients were grown overnight. 200 µl of helper and donor and 2 mL of recipient were separately washed with 10mM MgCl_2_ and then mixed together and washed again. The mix was then re-suspended in 30 µl of 10 mM MgCl_2_, plated on pre-warmed LB agar plates and incubated for 24-48 hours at 28˚C. Cells were then harvested, resuspended in 1ml of sterile 10 mM MgCl_2_ and plated on KB kanamycin nitrofurantoin plates. Selected mutants were screened for normal growth in KB, LB and M9.

### In vitro growth

Overnight cultures of *Psa* NZ13^kanR^ and *Psa* NZ45 were used to determine the *in vitro* growth of each strain in MGY alone or supplemented with 0.5 and 0.8mM CuSO_4_. 10mL liquid MGY cultures were established with a starting density of 10^5^ cfu mL^−1^ and shaken for up to three days. Bacterial growth was monitored by plating on KB kanamycin (*Psa* NZ13^kanR^), MGY 0.8mM CuSO_4_ (*Psa* NZ45). Three replicates per strain and media combination was used, and the experiment was repeated three times.

### *In planta* growth

Clonally propagated *Actinidia chinensis* var. ‘Hort16A’ plantlets were maintained at 20˚C with a light/dark period of 14/10 hours, 70% constant humidity. *Psa* NZ13^kanR^ and *Psa* NZ45 were grown on KB agar plates for 48h at 28˚C. Inoculum with a final optical density (OD_600_) of 0.2 of either strain was prepared in 50 ml 10mM MgCl_2_ with 0.002% of Silwet. Three to four week old plantlets were inoculated by dipping the aerial parts in the inoculums solution for 5 seconds. Five separate plantlets were dip inoculated for each treatment. For experiments assessing *in planta* growth in copper-sprayed plantlets, Nordox75 was used at the recommended dosage of 0.375g L^−1^.

(www.kvh.org.nz/spray_products). Dip-inoculated plantlets were allowed to dry, then sprayed adaxially and abaxially with Nordox75 until runoff to ensure complete coverage. Bacterial growth was monitored 0, 3 and 7 days post inoculation. 1 cm^2^ disk leaves were cut using a sterile cork borer, surface sterilized in 70% ethanol and ground in 200 µl 10mM MgCl_2_. Serial dilutions of the homogenate were plated on KB kanamycin to count *Psa* NZ13^kanR^ and MGY 0.8mM CuSO_4_ to count *Psa* NZ45.Each experiment was repeated 3 times.

### *In vitro* and *in planta* competition assays

*In vitro* and *in planta* competition assays were conducted as described earlier for single strains, except that *Psa* NZ45 and *Psa* NZ13^kanR^ were coinoculated in a 1:1 mix. Bacterial growth was monitored by plating serial dilutions on KB kanamycin (*Psa* NZ13^kanR^), MGY 0.8mM CuSO_4_ (*Psa* NZ45) and on MGY kanamycin 0.8mM CuSO_4_ (*Psa*NZ13^kanR^ that acquired copper resistance). *In vitro* assays had three replicates per strain, *in planta* assays were conducted using five replicates, each experiment was repeated three times. Fitness was calculated as ratio between their Malthusian Parameters (Lenski *et* al., 1991).

### ICE integration screening

Primers used in this study are listed in Supp Table 2. Four primers were designed to detect the genomic location of ICE integration: two specific for the integrases at the end of each ICEs (*IntPsaNZ45*, *IntPsaNZ13*) and two for the ICE insertion site on the chromosome, annealing to the *clpB* (*att-1* site) and *queC* (*att-2* site) genes. The primer combination (*IntPsaNZ45-att-2*, *IntPsaNZ45-att-1, IntPsaNZ13-att-1*, and *IntPsaNZ13-att-2*) indicates the location of the ICEs. Another two sets of primers were designed to amplify either CuR (*copA*) or enolase genes present in the VR of *Psa*_*NZ45*_ICE_Cu and *Psa*_NZ13_ICE_eno, respectively. PCRs were performed using Thermo Scientific *Taq DNA Polymerase* following the manufacturer’s instructions.

### ICE mobilization assay

*Psa* NZ45 was used as the ICE donor. Strains tested in the mobilization assays included are listed in order of divergence relative to the donor: *Psa* K28 (biovar 2) (McCann *et al*., 2013), *Psa* J31 (biovar 1) (McCann *et al*., 2013), *Peusomonas syringae* pv. *actinidifoliorum* NZ9 (McCann *et al*., 2013), *Pseudomonas syringae* pv. *tomato* DC3000 (Buell *et al*., 2003), *Pseudomonas syringae* pv *phaseolicola* 1448a (Teverson, 1991), *Pseudomonas syringae* H24 and H33 (isolated from kiwifruit; C. Straub, unpublished data), *Pseudomonas fluorescens* SBW25 (Zhang *et al*., 2006) and *Pseudomonas aeroginosa* PAO1 (Holloway, 1955). The copper sulphate MIC was determined for all tested recipient strains, which were all tagged with kanamycin Tn*5.* A biparental mating was performed using 2 mL and 200µl of washed recipient and *Psa* NZ45 cells, respectively. The cells were mixed, centrifuged briefly and resuspended in 30µl of 10 mM MgCl_2_ alone, 10 mM MgCl_2_ with 0.5mM CuSO_4_ or 30 µl of 1 cm^2^ kiwifruit plantlet macerate in 200µl of 10 mM MgCl_2_ if requested. The cell mixture was plated onto solid media (M9 plates) and incubated at 28˚C for 48 hours. Cells were then harvested and resuspended in 1ml of sterile 10 mM MgCl_2_. Serial dilutions were plated on KB kanamycin to count the total number of recipients and on MGY amended with kanamycin and copper sulphate at recipient MIC to count transconjugants.

### Biosecurity and approval

All worked was performed in approved facilities and in accord with APP201675, APP201730, APP202231.

## ACKNOWLEDGMENTS

We gratefully acknowledge Zespri International Limited and Te Puke Fruit Growers Association for financial support. The sponsors had no role in the design, collation, or interpretation of data. We thank kiwifruit growers in Te Puke for the access to orchards, Denis Robinson for providing Nordox75, and Daniel Rexin for assistance with isolating *Psa* from kiwifruit leaves.

## SUPPORTING INFORMATION

**Figure S1.**
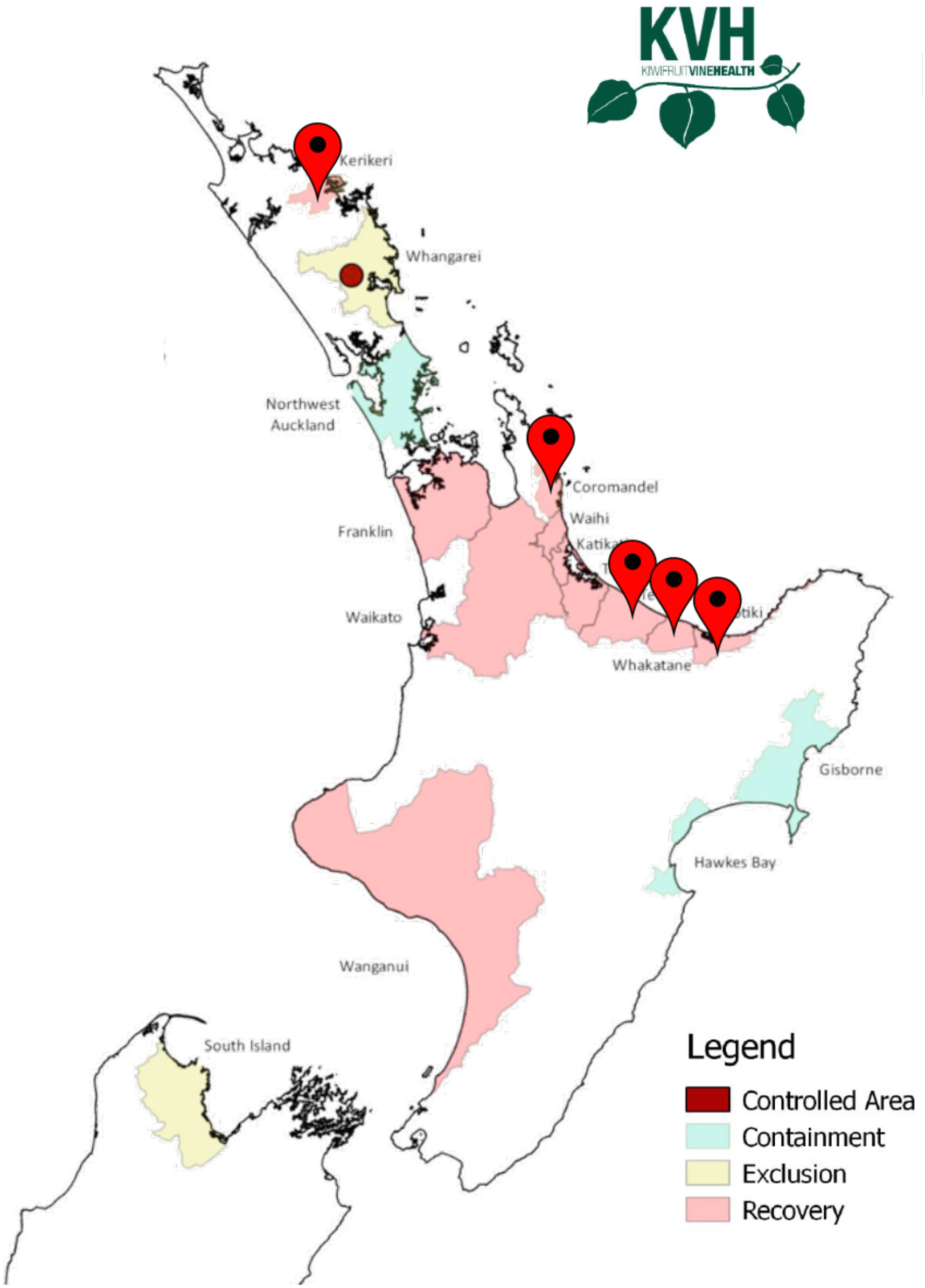
New Zealand kiwifruit growing regions with isolation sites of copper resistant *Psa.* Map was modified from regional classification map of June 2016 (Kiwi Vine Health (KVH)).

**Figure S2.**
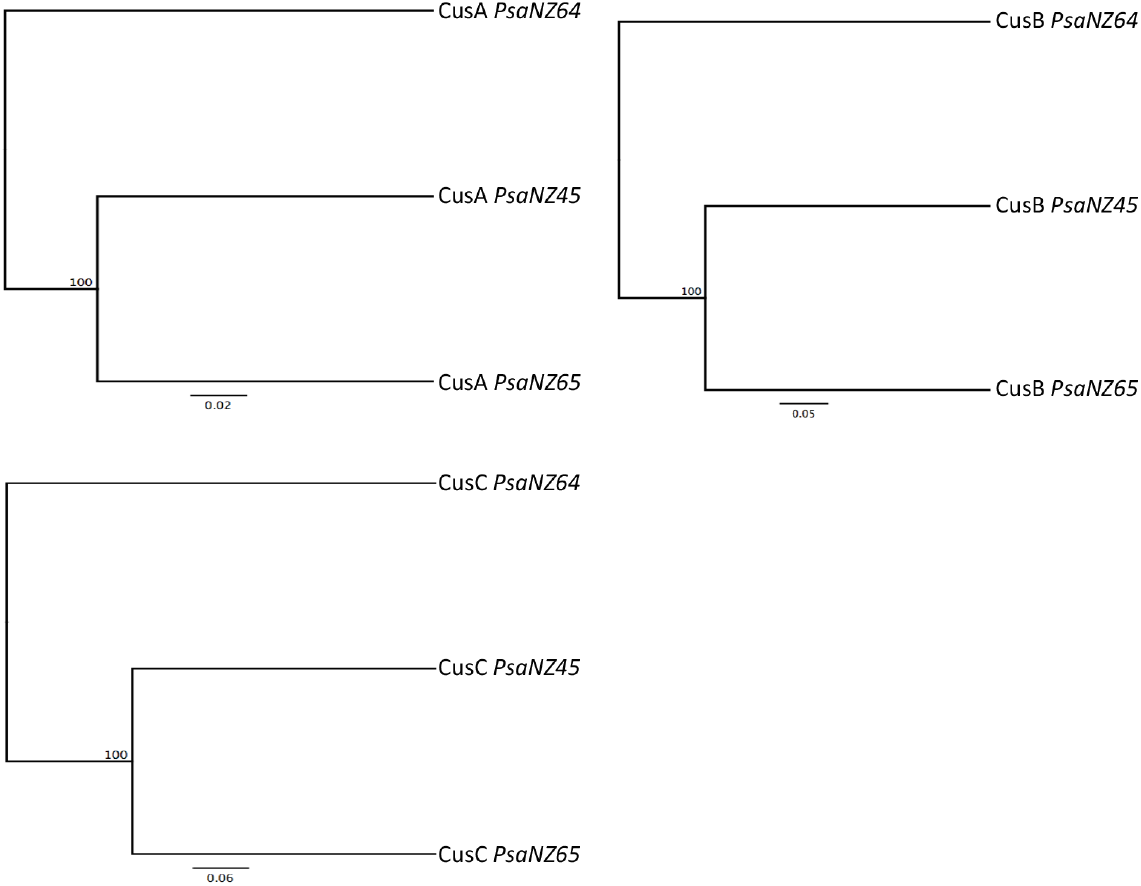
UPMGA tree of the Cus system proteins in *Psa* NZ. Bootstrap values are shown at each node.

**Figure S3.**
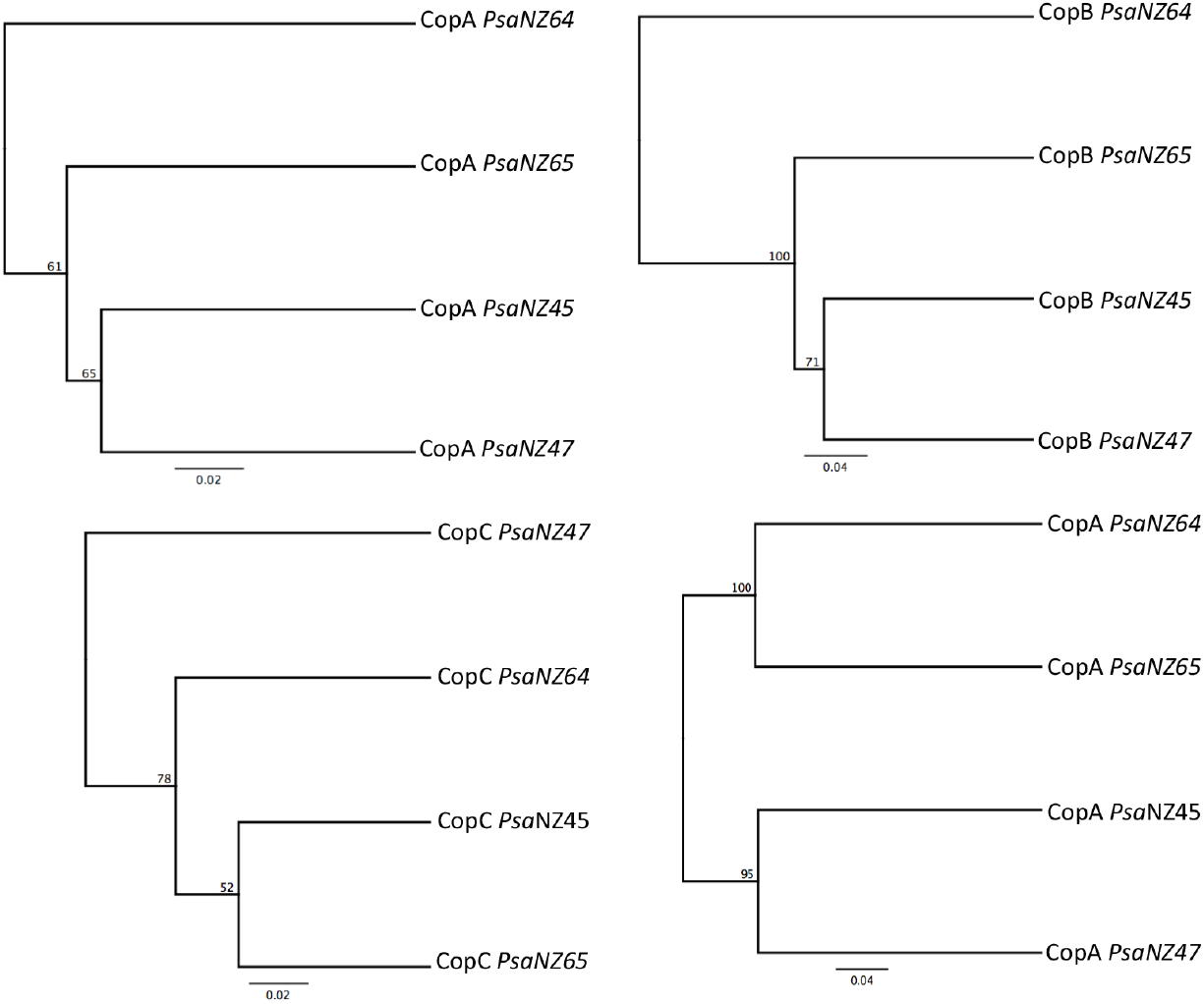
UPMGA trees of the Cop proteins genes in *Psa* NZ. Bootstrap values are shown at each node.

**Figure S4.**
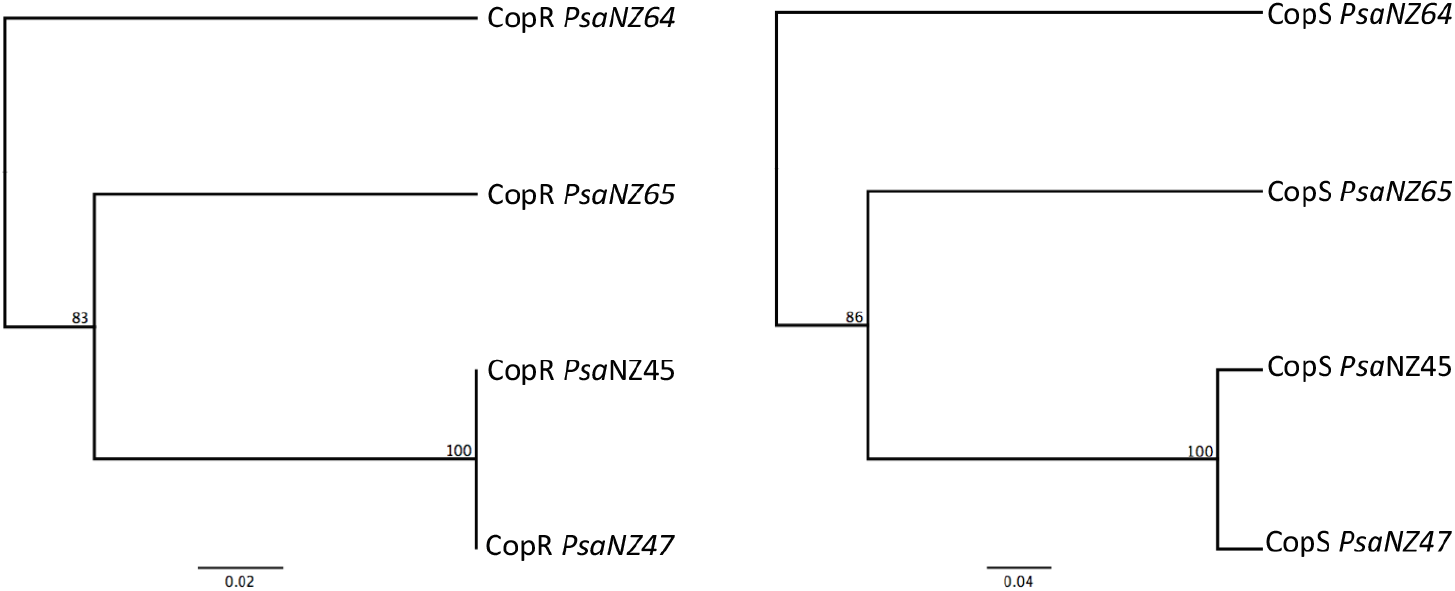
UPMGA trees of CopR and CopS proteins in *Psa* NZ. Bootstrap values are shown at each node.

**Figure S5.**
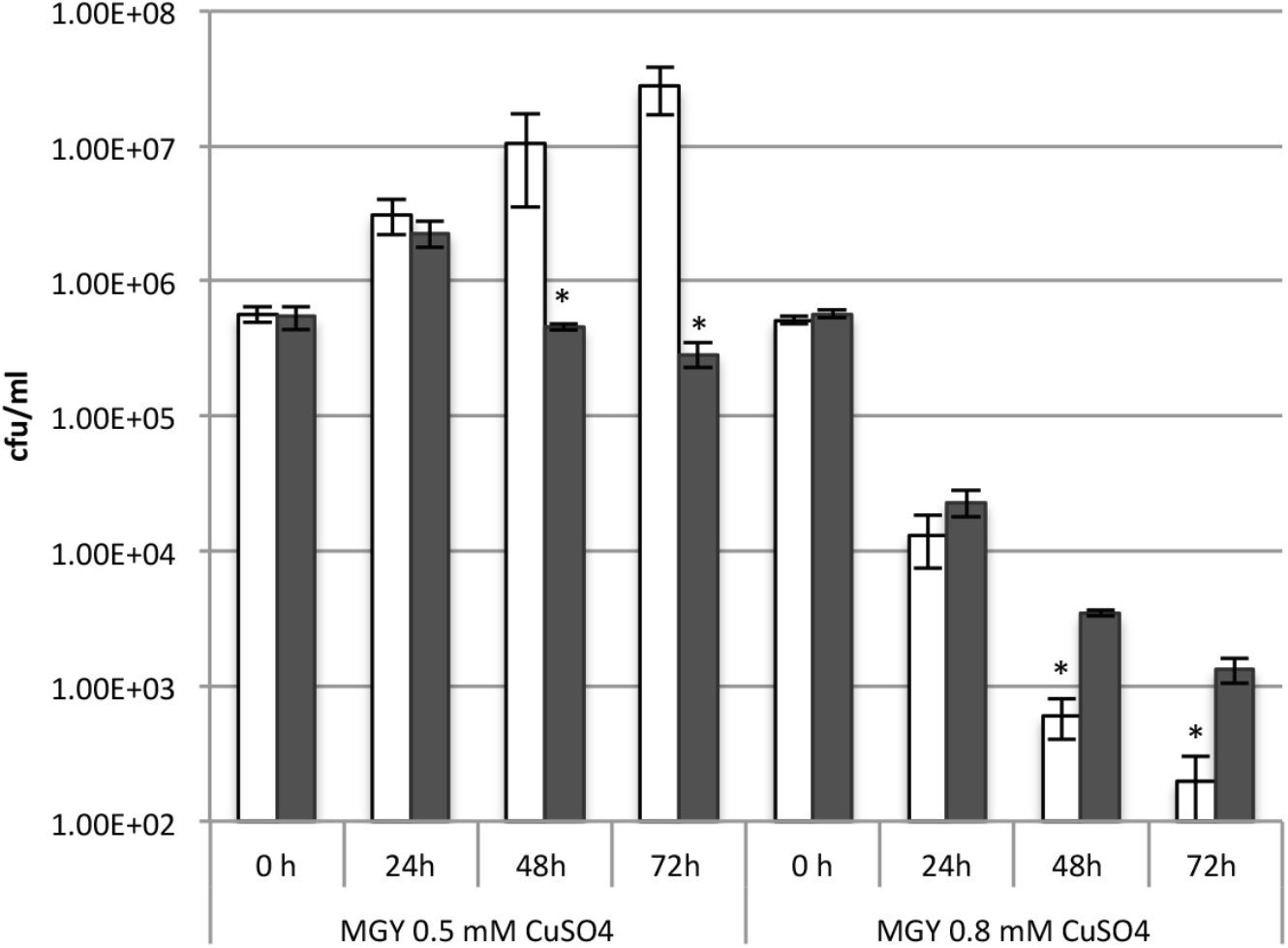
Density of single and co-cultured *Psa* in liquid MGY supplemented with 0.5 m and 0.8 mM CuSO_4_. *Psa* NZ13 was cultured alone (white bars) or co-cultured with *Psa* NZ45 (grey bars). Data are means and standard deviation of three independent cultures. *indicates significance at 5% level by one-tailed *t*-test.

**Figure S6.**
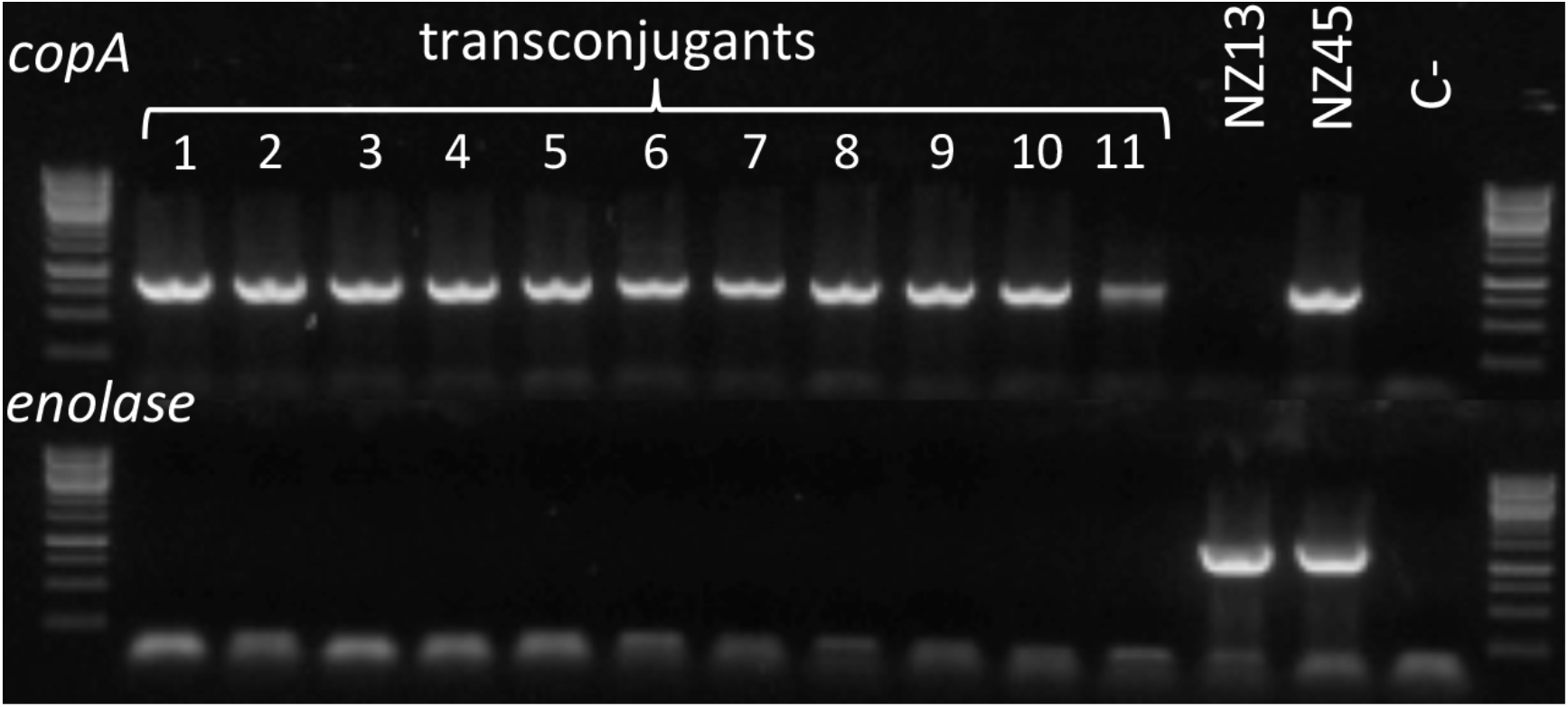
Analysis of the presence of the variable region (VR) of 948 *Psa*_NZ45_ ICE_Cu and *Psa*_NZ13_ ICE_eno in 11 *Psa* NZ13 transconjugants. PCRs were carried out to detect *copA* (VR of *Psa*_NZ45_ICE_Cu) or *enolase* genes (VR of 950 *Psa*_NZ13_ICE_eno). Controls of *Psa* NZ13 and *Psa* NZ45 show one and two bands, indiciative of *Psa*_NZ13_ICE_eno in *Psa* NZ13 and both *Psa*_NZ13_ICE_eno*Psa*_NZ45_ICE_Cu 952 and in *Psa* NZ45, respectively. All transconjugants, lanes 1-11 have acquired *Psa*_NZ45_ICE_Cu.

**Figure S7.**
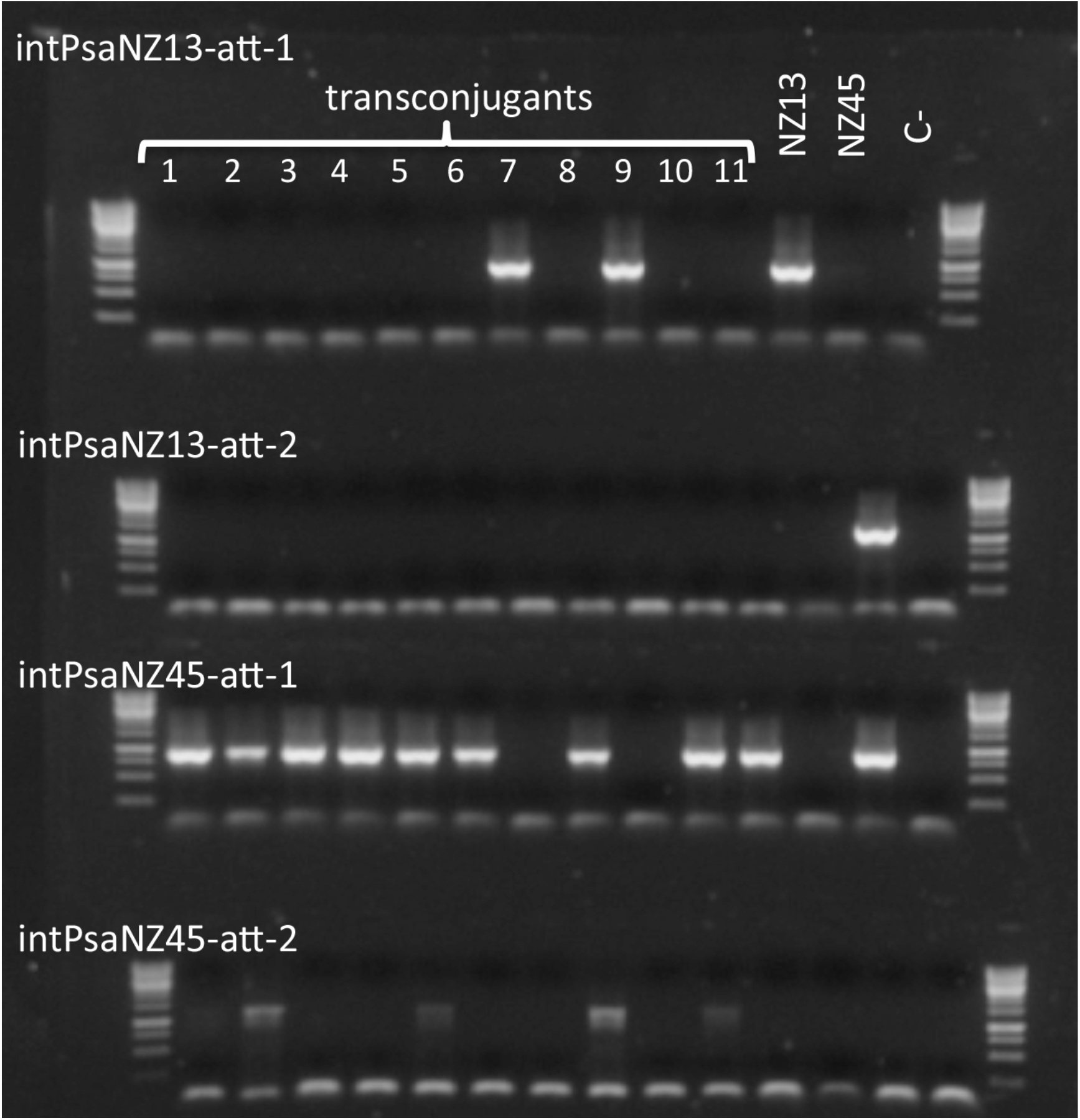
Analysis of the insertion site of the *Psa*_NZ13_ICE_eno and *Psa*_NZ45_ICE_Cu in 11 *Psa* NZ13 transconjugants. PCRs were to detect the integration of *Psa*_NZ13_ICE_eno in the *att-1* or *att-2* sites (*intPsaNZ13-att-1* and 968 *intPsaNZ13-att-2*) and the integration of *Psa*_NZ45_ICE_Cu in the *att-1* or *att-2* sites (*intPsaNZ45-att-1* and *intPsaNZ45-att-2*). Controls of *Psa* NZ13 and *Psa* NZ45 show that in *Psa* NZ13 the *Psa*_NZ13_ICE_eno is integrated in the *att-1* site and in *Psa* NZ45 the *Psa*_NZ13_ICE_eno is integrated in the *att-1* and the *Psa*_NZ45_ICE_Cu in the *att-2* site.

**Table S1.** *In planta* transfer of *Psa*_NZ45_ICE_cu from *Psa* NZ45 to *Psa* NZ13 at different founding ratios of donor and recipient. Donor and recipient strains were dip-inoculated onto Hort16A leaves at different founding ratios and frequency of recipients determined at days 3 and 7

|  | Frequency of *Psa*<sub>NZ45</sub>ICE_Cu transconjugants |  |
| --- | --- | --- |
| *Psa* NZ13 : *Psa* NZ45 | day 3 | day 7 |
| 1 : 1 | $(2.05 \pm 0.63)^{-2}$ | $(2.28 \pm 0.7)^{-2}$ |
| 1 : 0.1 | $(1.34 \pm 0.52)^{-2}$ | $(3.14 \pm 2.3)^{-2}$ |
| 0.1 : 1 | $(1.68 \pm 0.68)^{-2}$ | $(1.88 \pm 0.81)^{-2}$ |

**Table S2.**
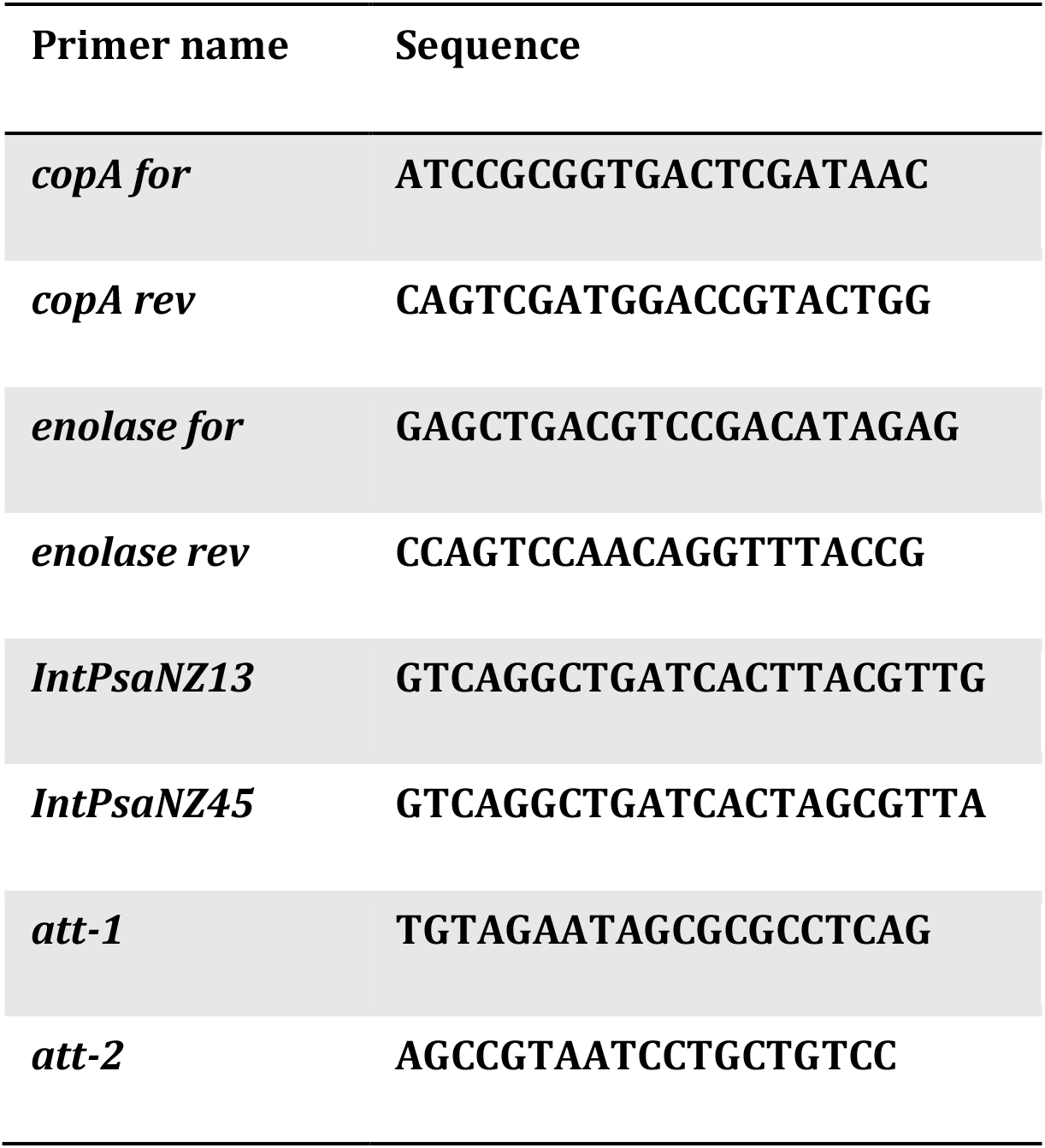
Primers used for *Psa*_NZ13_ICE_eno and *Psa*_NZ45_ICE_Cu detection and integration loci.

